# Lasting consequences on physiology and social behavior following cesarean delivery in prairie voles

**DOI:** 10.1101/2022.05.22.492927

**Authors:** William Kenkel, Marcy Kingsbury, John Reinhart, Murat Cetinbas, Ruslan I. Sadreyev, C. Sue Carter, Allison Perkeybile

## Abstract

Cesarean delivery is associated with diminished plasma levels of several ‘birth-signaling’ hormones, such as oxytocin and vasopressin. These same hormones have been previously shown to exert organizational effects when acting in early life. For example, our previous work found a broadly gregarious phenotype in prairie voles exposed to oxytocin at birth. Meanwhile, cesarean delivery has been previously associated with changes in social behavior and metabolic processes related to oxytocin and vasopressin. In the present study, we investigated the long-term neurodevelopmental consequences of cesarean delivery in prairie voles. After cross-fostering, vole pups delivered either via cesarean or vaginal delivery were studied throughout development. Cesarean-delivered pups responded to isolation differently in terms of their vocalizations (albeit in opposite directions in the two experiments), huddled in less cohesive groups under warmed conditions, and shed less heat. As young adults, we observed no differences in anxiety-like or alloparental behavior. However, in adulthood, cesarean-delivered voles of both sexes failed to form partner preferences with opposite sex conspecifics. In a follow-up study, we replicated this deficit in partner-preference formation among cesarean-delivered voles and were able to normalize pair-bonding behavior by treating cesarean-delivered vole pups with oxytocin (0.25 mg/kg) at delivery. Finally, we detected minor differences in regional oxytocin receptor expression within the brains of cesarean-delivered voles, as well as microbial composition of the gut. Gene expression changes in the gut epithelium indicated that cesarean-delivered male voles have altered gut development. These results speak to the possibility of unintended developmental consequences of cesarean delivery, which currently accounts for 32.9% of deliveries in the U.S. and suggest that further research should be directed at whether hormone replacement at delivery influences behavioral outcomes in later life.

## INTRODUCTION

In recent decades, the rate of delivery by cesarean section (CS) has risen to 32.9% of all births in the United States (Boerma et al., 2018; Martin et al., 2019; Zhang et al., 2019). Despite its widespread adoption, we have only recently begun to explore the possibility that CS delivery could produce neurodevelopmental consequences (Hyde et al., 2012), spurred on by epidemiological associations between CS delivery and increased risk profiles for autism spectrum disorders, attention deficit / hyperactivity disorder, and overweight / obesity (Cai et al., 2018; Curran et al., 2015b; Hansen et al., 2018; Masukume et al., 2019; Mueller et al., 2019; Sitarik et al., 2020; Yip et al., 2017; Zhang et al., 2019). While recent analyses suggest the association between CS and neurodevelopmental disorders arises from unmeasured familial confounding (Curran et al., 2015a; Zhang et al., 2021), other subclinical outcomes may still occur. Furthermore, purely correlational studies in humans remain unable to resolve this question definitively, and experimental studies in humans are ethically infeasible. This leaves experimental studies in animal models to address this gap in our knowledge. Whereas humans and rodents differ in terms of cortical development at birth (Semple et al., 2013), all mammals must navigate a common set of challenges pertaining to the initiation of independent homeostasis upon delivery. It is here that two potential mechanisms for how CS could impact development arise. First, CS may impact development through disturbances of microbial colonization that typically begins upon passage through the birth canal (Montoya-Williams et al., 2018; Torres-Fuentes et al., 2017) and is known to influence gut-brain signaling (Moya-Perez et al., 2017). Secondly, CS may affect development through diminished levels of the ‘birth-signaling hormones’ during a sensitive period in development.

Maternal and fetal levels of several ‘birth-signaling hormones’, such as oxytocin (OXT) (Kuwabara et al., 1987; Marchini et al., 1988), vasopressin (Burckhardt et al., 2014; Wellmann et al., 2010), glucocorticoids, epinephrine, and norepinephrine all surge during vaginal delivery (VD) (Elbay et al., 2016; Lagercrantz and Slotkin, 1986; Talbert et al., 1977; Vogl et al., 2006). Levels of these hormones in the plasma of the neonate are lower after CS delivery compared to those of VD neonates, particularly if CS occurs prior to the onset of labor (Kenkel, 2020). Furthermore, CS delivery is often performed relatively early in labor, prior to the recommendations laid out by the American College of Obstetricians and Gynecologists (Zhang et al., 2010), resulting in even lower birth signaling hormone levels (Lagercrantz and Slotkin, 1986; Pohjavuori and Fyhrquist, n.d.). The surging levels of the ‘birth-signaling hormones’ are important because they help accomplish the transition to independent homeostasis that must begin at birth (Kenkel, 2020; Kenkel et al., 2014). Lower levels of these hormones raise the prospect of developmental programming because the perinatal period is a sensitive period in development (Hammock, 2015; Kenkel, 2020; Kenkel et al., 2014; Miller and Caldwell, 2015). One domain that appears particularly sensitive to early-life OXT manipulation is social behavior (Bales et al., 2007b; Bales and Carter, 2003a, 2003b; Kenkel et al., 2019b). Most studies of CS in humans have focused on clinical outcomes, such as autism spectrum disorder, and there have also been findings to suggest CS impacts specific aspects of human social behavior as well. A study of ~11,000 children found CS delivery associated with a delay in personal social skills at 9 months (Al Khalaf et al., 2015), and in a study of early childhood, emergency CS was associated with peer problems (Rutayisire et al., 2016).

In the present study we chose to explore the neurodevelopmental consequences of CS delivery with a focus on social behavior (Experiment 1) and, in a follow-up study (Experiment 2), we attempted to normalize development through direct administration of OXT to CS offspring. Both experiments were heavily influenced by our prior work that examined how OXT exposure on the day of delivery influenced neurodevelopment in the prairie vole (Kenkel et al., 2019b). We selected a battery of tests aimed at behaviors known to be under the regulation by OXT, including the production of isolation-induced ultrasonic vocalizations (USVs) on postnatal days (PNDs) 1 and 4 (Kenkel et al., 2019b), behavioral thermoregulation on PND 7 (Harshaw et al., 2018), consolation behavior toward a stressed cage-mate on PND 45-46 (Burkett et al., 2016), anxiety-like behavior on PND 48-52 (Klenerova et al., 2009), alloparental responsiveness on PND 50-55 (Kenkel et al., 2019b), and pair-bonding on PND 60-66 (Kenkel et al., 2019b). Upon completion of this behavioral battery, subjects were sacrificed and brain tissue collected for autoradiographic analysis of the OXT receptor (OTR) and vasopressin receptor (V1aR), as well as gut tissue and cecal contents, which were analyzed for histology and microbiome.

## METHODS

### Subjects

Subjects were laboratory-bred prairie voles (*Microtus ochrogaster*), descendants of a wild-caught stock captured near Champaign, Illinois. The stock was systematically outbred. Breeding pairs were housed in large polycarbonate cages (44cm x 22cm x 16cm) and same sex offspring pairs were housed in smaller polycarbonate cages (27cm x 16cm x 16cm) after weaning on PND 20. Animals were given food (high-fiber Purina rabbit chow) and water ad libitum, with enviro-dry and cotton nestlets for nesting material in breeding cages, and were maintained at room temperature (22°) on a 14:10 light:dark cycle. To generate experimental subjects, pairs were created using stud males aged 60-240 days and primiparous adult female voles aged 60-90 days. To minimize the contributions of prematurity, we developed a timed mating paradigm to ensure accurate prediction of expected delivery. Female prairie voles undergo induced estrus and induced ovulation, which led us to carefully control the timing and duration of access to males. First, adult females were introduced to adult males in a novel cage into which we introduced dirty bedding from the male’s home cage to induce estrus (Carter, 1987). Males and females cohabited without mating for 24 hours, at which point a perforated cage divider was introduced, separating male from female. This cage divider remained for ~72 hours, at which point it was removed and mating confirmed by observation. Females were considered term pregnant if they gained >20g body weight and were 21.5 days past an observed mating.

Pregnant female dams bore litters either via vaginal delivery (VD) or cesarean section (CS) approximately 21.5 days after mating. CS deliveries typically occurred on the same day as VD deliveries and while there is some variability in delivery time, this CS paradigm typically achieved delivery ~12 hours prior to expected delivery. In the CS condition, surgical delivery occurred via laparotomy following 90 seconds of CO2 anesthesia according to the protocol developed by the Forger lab (Castillo-Ruiz et al., 2018; Hoffiz et al., 2021), an approach which avoids the confound of pharmacological anesthesia. CS delivery was thus a terminal procedure so as to avoid the confound of surgery affecting maternal behavior and because CS vole dams do not respond maternally to pups (Hayes and De Vries, 2007). Following delivery (CS) or discovery within 16 hours of delivery (VD), litters were cross-fostered as same-condition litters to parents that had had litters of their own within the previous 3 days. In Experiment 2, we added a third condition: CS+OXT in which CS-delivered pups were immediately injected subcutaneously with 0.25 mg/kg OXT (Pitocin). Likewise, in Experiment 2, CS and VD pups were injected subcutaneously with saline immediately after delivery or discovery. While endogenous OXT levels at delivery remain unknown in this species, we selected this dose based on our previous work (Kenkel et al., 2019b), where 0.25 mg/kg OXT was not sufficient to trigger labor in term pregnant female prairie voles. This dose is also comparable to previous work aimed at averting the consequences of CS by means of OXT administration (0.1 – 1 mg/kg (Boksa et al., 2015; Morais et al., 2021)).

Offspring were then studied throughout development according to the following protocols. For Experiment 1, a total of 23 CS and 23 VD litters were generated, and for experiment 2, 8 CS, 8 VD, and 7 CS+OXT litters were generated. However, not all offspring were used for all measures, owing either to logistic challenges or data loss due to experimenter error. Final sample sizes for each measure are specified below. All procedures were carried out in accordance with the National Institutes of Health Guide for the Care and Use of Laboratory Animals and with the approval of the Bloomington Institutional Animal Care and Use Committee (BIACUC).

### Ultrasonic vocalizations (USVs)

According to previously published methods (Kenkel et al., 2019b), we recorded isolation-induced USVs from each of a litter’s pups on PNDs 1 and 4. Briefly, pups were removed from the nest and tested individually for 5 minutes at room temperature. USVs were recorded using an Avisoft UltraSoundGate 116Hme microphone, sampling at 192 KHz. Vocalizations were detected using Avisoft-SASlab Pro version 5.2.07. A total of 68 CS pups and 88 VD pups had USVs recorded for Experiment 1, and 32 CS, 33 VD pups and 25 CS+OXT pups had USVs recorded for Experiment 2.

### Thermography

On PND-7, offspring were tested as litters to characterize thermoregulation in terms of skin surface temperature and behavior. In Experiment 1, a total of 19 CS and 21 VD litters were tested. However, a combination of experimenter error and technical issues prevented us from completing a sufficient number of litters to conduct this analysis in Experiment 2. Litters were removed from the nest and placed into a jacketed beaker wherein ambient temperature could be manipulated via passage of heated / chilled water through the walls of the chamber. For 30 minutes litters were exposed to 33° (warm phase), at which point the chamber’s water was changed to 22°C and once the chamber walls equilibrated to this new temperature (~2-4 minutes), observations resumed for another 30 minutes (cool phase). During observations, measures were collected once per minute using both webcam images and thermographic infrared images captured by means of an ICI 9640-P infrared camera (Infrared Cameras Inc.; ICI; Beaumont, TX). The resulting images were then analyzed using a custom Matlab code. For the behavioral measures (webcam), images were binarized and litter huddles analyzed in terms of: 1) the total number of discontinuous clumps and/or lone pups, and 2) the total perimeter of all clumps. Thus, a more cohesive litter would have fewer clumps and a shorter perimeter. For the thermographic measures (infrared), images were analyzed in terms of the number of pixels > 30.5°C. The parameter of number of pixels > 30.5°C was chosen because it best correlated with manually collected intrascapular measures of surface temperature in mouse pups (Harshaw et al., manuscript in preparation). While the walls of the chamber were 33°C during the warm phase, the floor of the chamber consisted of a thin foam sheet which remained at < 30°C. Thermographic images were analyzed in terms of the number of pixels per image over 30.5°C using a repeated measures linear mixed effects model with birth mode (CS vs. VD), time as main effects and accounting for litter weight as a random effect.

### Consolation

Between PND-45 and 46, a maximum of one male and one female from each litter were tested for conciliatory behavior exhibited toward a stressed same-sex cage mate, typically a sibling. In each pair of animals, one animal was subjected to 30 minutes of restraint stress and the behavior of the remaining animal was then assessed for 30 minutes when the stressed animal was returned to the cage. Behavior was analyzed by two observers blind to experimental condition and also using idTracker. Manually scored conciliatory behaviors included: allogrooming, huddling and sniffing. Automatically scored behaviors included: average distance between the two animals and the time spent within one body length of the stressed animal. The consolation test was only included in Experiment 1, for which a total of 12 CS males, 5 CS females, 8 VD males and 8 VD females were included.

### Open Field

Open Field testing was carried out according to previously used methods (Kenkel et al., 2019b). Between PND-48 and 52, a maximum of one male and one female from each litter were tested individually in a plexiglass arena (42 x4 2 cm) for 20 minutes. Behavior was analyzed using idTracker (Perez-Escudero et al., 2014) in terms of total distance traveled and time spent in the center of the arena. For Experiment 1, a total of 16 CS males, 7 CS females, 17 VD males and 20 VD females were tested; for Experiment 2, a total of 13 CS males, 21 CS females, 10 VD males, 20 VD females, 11 CS+OXT males, and 13 CS+OXT females were tested.

### Alloparenting

Testing of alloparental responsiveness was carried out according to previously used methods (Kenkel et al., 2019b). Between PND-50 and 55, a maximum of one male and one female from each litter were tested individually in a novel cage for 20 minutes for caregiving behavior directed toward a novel, unrelated pup aged 1-3 days. Measures of interest included time spent: Not in contact with the pup (‘Not in Contact’), Sniffing or making incidental contact with the pup (‘Contact’), Licking and/or grooming the pup (‘Licking / Grooming’), and huddling over the pup (‘Huddling’). In Experiment 1, a total of 17 CS males, 9 CS females, 20 VD males and 23 VD females were tested; in Experiment 2, a total of 11 CS males, 18 CS females, 8 VD males, 19 VD females, 9 CS+OXT males, and 13 CS+OXT females were tested. Behavior was scored using BORIS, an open-source event-logging software (“BORIS: a free, versatile open source event logging software for video/audio coding and live observations - Friard - 2016 - Methods in Ecology and Evolution - Wiley Online Library,” n.d.) by an experimentally blind observer.

### Pair-bonding and Partner Preference Testing (PPT)

Testing of pair-bonding and partner preference formation was carried out according to previously used methods (Kenkel et al., 2019a). Between PND-60 and 66, a maximum of one male and one female from each litter were paired with an opposite sex adult and allowed to cohabit for 24 hours, after which time subjects were tested in the PPT (Figure 4). Subjects were presented with the familiar opposite-sex animal they had cohabited with (‘Partner’) as well as a novel opposite-sex animal (‘Stranger’), each of whom were tethered in individual chambers while the subject was able to freely travel. Behavior was analyzed using idTracker (Kenkel et al., 2019a; Perez-Escudero et al., 2014) in terms of the time subjects spent in close proximity to either the Partner or Stranger. In addition to proximity, we applied a further immobility criterion to better correspond with the ‘side by side contact’ measure of the traditional, manually-scored PPT (Kenkel et al., 2019a). In Experiment 1, a total of 17 CS males, 11 CS females, 22 VD males, and 23 VD females were tested; in Experiment 2, a total of 4 CS males, 13 CS females, 6 VD males, 9 VD females, 10 CS+OXT males and 10 CS+OXT females were tested.

**Figure 1.**
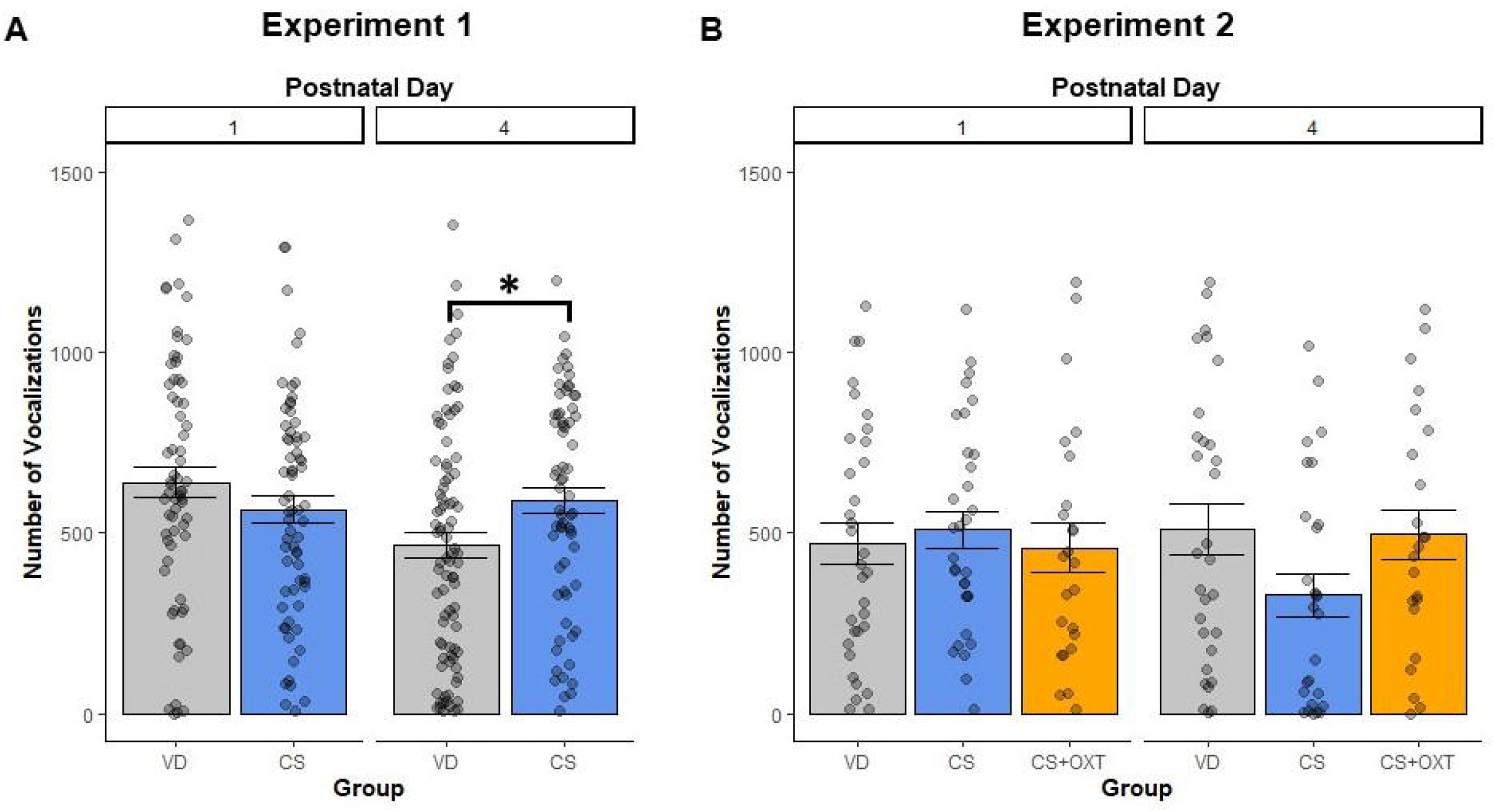
Production of ultrasonic vocalizations (USVs) by vole pups delivered either via cesarean section (CS) or vaginal delivery (VD). In Experiment 1 (panel A), CS pups produced significantly more USVs on postnatal day (PND) 4 and VD pups produced significantly fewer USVs on PND 4 compared to PND 1 (* denotes p = 0.018 and p = 0.002 respectively, n = 288 total recordings from 43 litters (n = 22 VD, n = 21 CS)). However, in Experiment 2 (panel B), CS pups tended to produce fewer USVs than VD pups on PND-4 (p = 0.061, n = 172 total recordings from 23 litters (n = 8 VD, n = 8 CS, n = 7 CS+OXT)).

**Figure 2.**
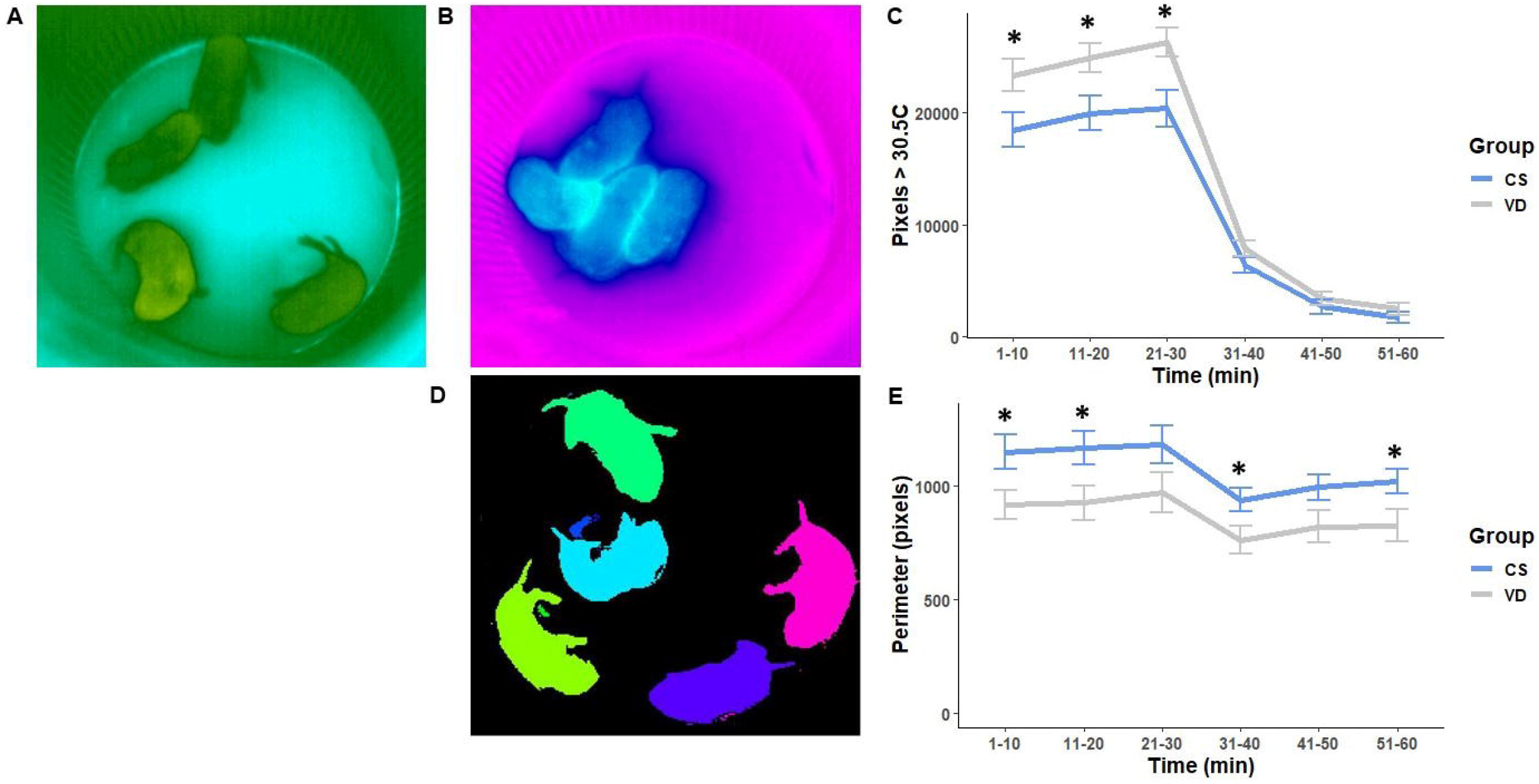
Thermoregulatory behavior in CS and VD vole litters on PND-7. Nineteen CS and 21 VD litters were tested for 30 minutes at 33°C followed by 30 minutes at 22°C. Representative thermographic images isare shown in panel A (warm) and B (cool). CS litters had less warm surface temperatures compared to VD litters during the warm phase (panel C, * denotes p < 0.025 for all comparisons). A representative image is shown in panel D having undergone automated identification of separate blobs (note that the seemingly disconnected hindleg in blue would have been excluded for having been too small). CS litters’ huddles were less cohesive throughout the majority of the testing session, during both the warm and cool phases (panel E, * denotes p < 0.041). Data represent mean ± SEM.

**Figure 3.**
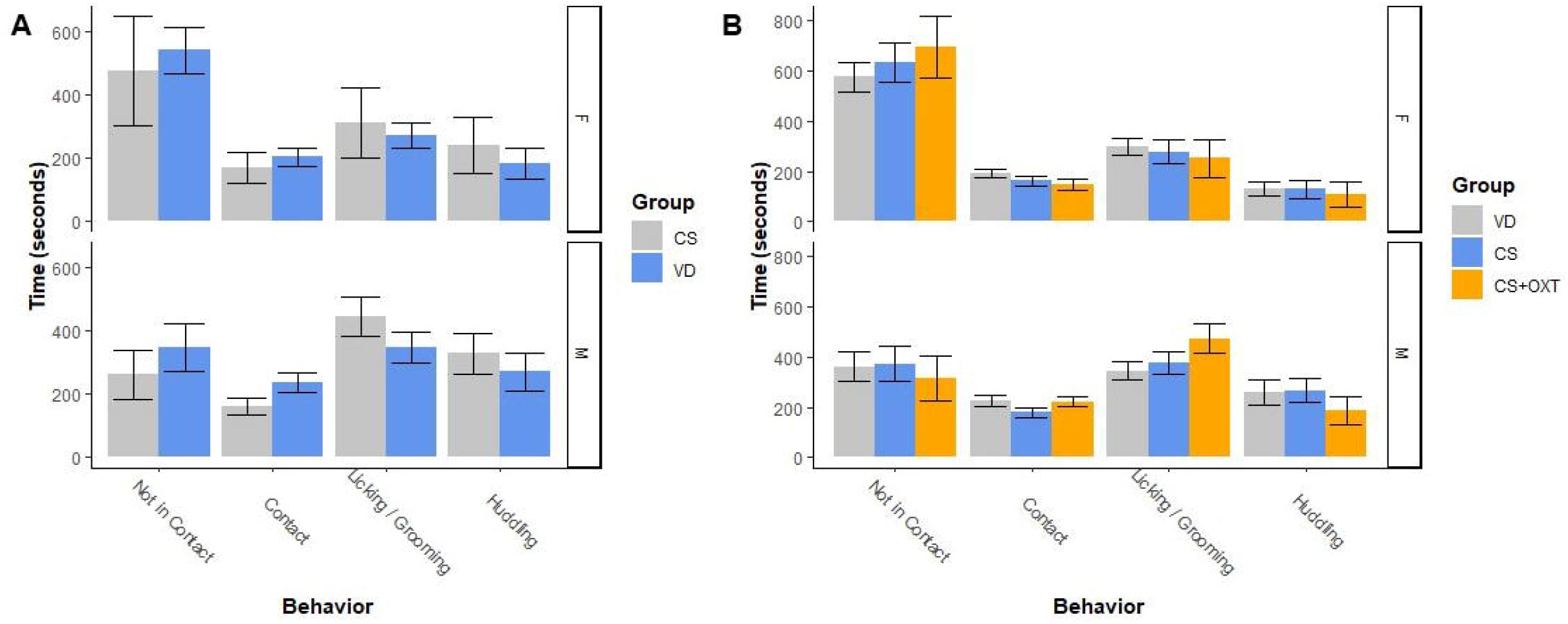
Alloparental responsiveness toward novel, unrelated pups. In neither Experiment 1 (A) nor Experiment 2 (B) did we detect a difference in alloparental behavior between CS and VD animals (p > 0.05 for all comparisons). In both experiments, males were found to be more alloparental than females, as has been observed previously. Data represent mean ± SEM.

**Figure 4.**
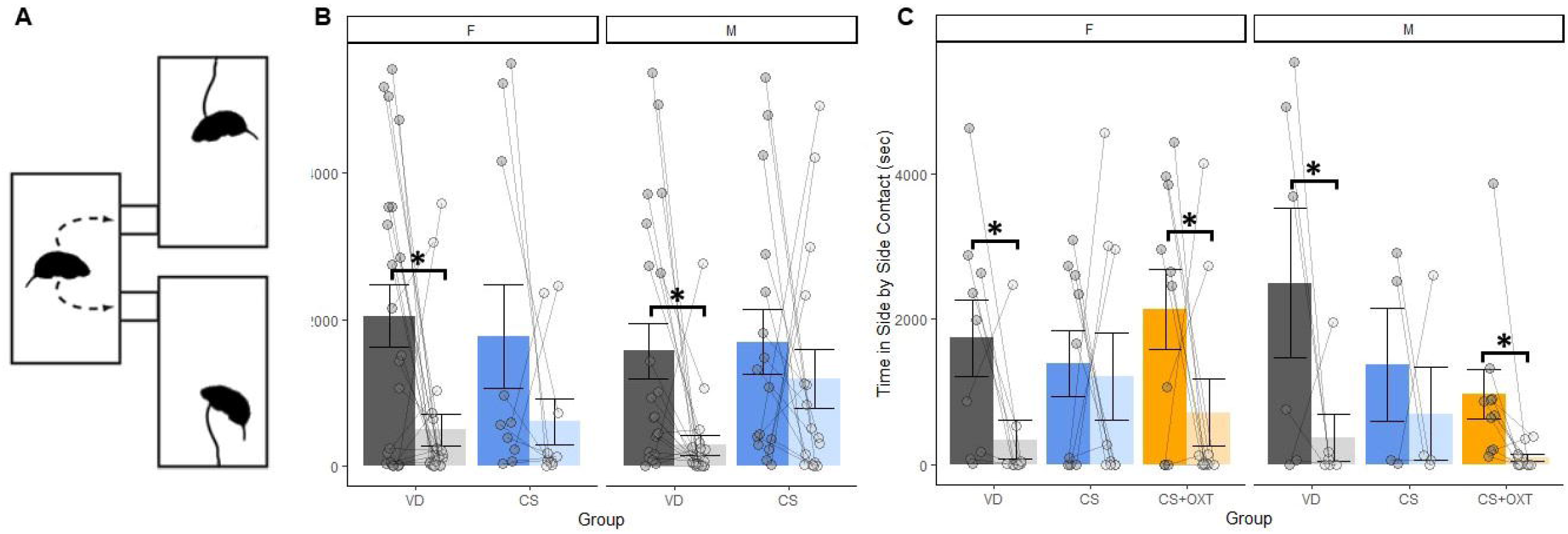
Partner preference formation in CS and VD voles on PND 60-68. The partner preference test (PPT) paradigm is shown in panel A. In Experiment 1, both female (F) and male (M) VD voles showed significant preferences for the Partner over the Stranger in terms of time spent in close social contact, as expected given prairie voles’ well-established social monogamy (panel B, * denotes p < 0.05 for both comparisons). Results from Experiment 1 include data from 17 CS males, 11 CS females, 22 VD males, and 23 VD females. In Experiment 2, VD voles again showed a significant partner preference, as did CS+OXT voles (panel C, * denotes p < 0.05 for all comparisons), whereas once again, CS animals showed no such preference. Results from Experiment 2 include data from 4 CS males, 13 CS females, 6 VD males, 9 VD females, 10 CS+OXT males and 10 CS+OXT females were tested. Data represent mean ± SEM.

### Sacrifice and brain tissue collection

In Experiment 1, one day after the completion of the PPT, a maximum of one male and one female from each litter were anesthetized with isoflurane and sacrificed for brain tissue collection. Brains were flash frozen on dry ice and section at 20μm for autoradiography according to previously published methods (Kenkel et al., 2019a, 2019b). Briefly, either the OXT receptor (OTR) or vasopressin receptor (V1aR) were visualized using specific radiolabeled ligands and radiosensitive film. For OTR binding [^125^I]-ornithine vasotocin analog [(^125^I)OVTA] [vasotocin, d(CH_2_)_5_[Tyr(Me)^2^, Thr^4^, Orn^8^, (^125^I)Tyr^9^-NH_2_]; 2200 Ci/mmol] was used (NEN Nuclear, Boston, MA, USA). For V1aR binding ^125^I-lin-vasopressin [^125^I-phe-nylacetyl-D-Tyr(ME)-Phe-Gln-Asn-Arg-Pro-Arg-Tyr-NH_2_] (NEN Nuclear) was used. Slides were exposed to Kodak BioMaxMR film (Kodak, Rochester, NY, USA) with ^125^I microscale standards (American Radiolabeled Chemicals, Inc., St., Louis, MO, USA). Slides were exposed for 168 h for OTR binding and 96 h for V1aR binding. Three coronal slices were measured for each animal at each of three landmarks aimed at capturing the following brain regions: 1) nucleus accumbens, 2) lateral septum and medial preoptic area, and 3) amygdala and hypothalamus. The complete list of brain regions investigated for OTR included the: agranular insular cortex, bed nucleus of the stria terminalis, central amygdala, cingulate cortex, claustrum, entorhinal cortex, lateral septum, nucleus accumbens core and shell, paraventricular nucleus of the hypothalamus, and parietal cortex. The complete list of brain regions investigated for V1aR included the: basolateral amygdala, bed nucleus of the stria terminalis, central amygdala, laterodorsal thalamus, medial cingulate cortex, paraventricular nucleus of the hypothalamus, preoptic area, retrosplenial cortex, and ventral pallidum. A total of 12 CS males, 12 CS females, 12 VD males and 12 VD females were included.

Images of slides were digitally scanned at 1800dpi. Three images of each subject, matched for anterior-posterior position, were then registered to one another using Photoshop (Adobe Systems Inc., San Jose, CA). A measure of non-specific binding (NSB) was also taken for each section from a gray matter region where minimal binding is detected. The NSB value was subtracted from the binding value for each section and a mean was then calculated for each section. Optical density was converted into decays per minute (DPM) using the i125 standards. DPM were then measured within each atlas-defined ROI using custom designed Matlab code and a slightly modified version of the prairie vole brain atlas we originally developed for magnetic resonance imaging (Yee et al., 2016). Results were calculated as regional averages of DPM (which corresponds to receptor density). Images were then consolidated into group composites for the purpose of visualizing group differences. Heat maps were generated in Matlab for each group-by-group comparison by subtracting one group composite from another (Supplemental Figure 1).

### Sacrifice and gut tissue/stool collection

In Experiment 1, when subjects were anesthetized with isoflurane and sacrificed for brain tissue collection, gut tissue and stool were also collected. Intestinal tissue from the terminal ileum and proximal colon was rapidly dissected in cold PBS and immediately flash frozen in isopentane/dry ice and stored at −80° Celsius for RNA isolation and qPCR. Stool from the cecum was collected and flash frozen in isopentane/dry ice and stored at −80°C for microbiome analyses.

### RNA Extraction, cDNA synthesis, and qPCR

To extract RNA, intestinal tissue was homogenized in Trizol® (Thermo-Fisher Scientific), vortexed for 10 min at 2000 rpm and placed at room temperature (RT) for 15 min. Chloroform was then added to the Trizol solution at a dilution of 1:5 and vortexed for 2 min at 2000 rpm. Tissue samples sat at RT for 3 min, were centrifuged at 11,800 rpm at 4°C for 15 min and the aqueous phase was removed. Isopropanol was added to the aqueous phase at a dilution of 1:1 to precipitate RNA and samples were vortexed again. Samples sat at RT for 10 min, were centrifuged at 11,800 rpm at 4°C for 15 min and RNA pellets were rinsed two times in ice-cold ethanol (75%). Pellets were then resuspended in 20 μl of nuclease-free water and frozen at −80°C until cDNA synthesis. The QuantiTect Reverse Transcription Kit (Quiagen) was used to synthesize cDNA. For each tissue sample, 200ng of RNA in 12 μl nuclease-free H_2_O was pre-treated with gDNase at 42°C for 2 min. 8 μl of master (primer-mix plus reverse transcriptase) was then added to each sample. Samples were heated in a thermocycler to 42°C for 30 min followed by 95°C for 3 min. qPCR was run on a Mastercycler ep realplex (Eppendorf) using the SYBR Green PCR Kit (Quiagen). PCR primers for prairie vole genes were designed in the lab using NIH primer blast and NIH nucleotide search, Beacon Designer and Eurofins Oligo analyses tool. Primers were purchased from Integrated DNA technologies. See Table 1 for primer sequences. Relative gene expression was calculated using the 2-Δ△CT method, relative to 18 S (house-keeping gene) and the lowest sample on the plate (Williamson et al., 2011; Livak & Schmittgen, 2001).

**Table 1.**
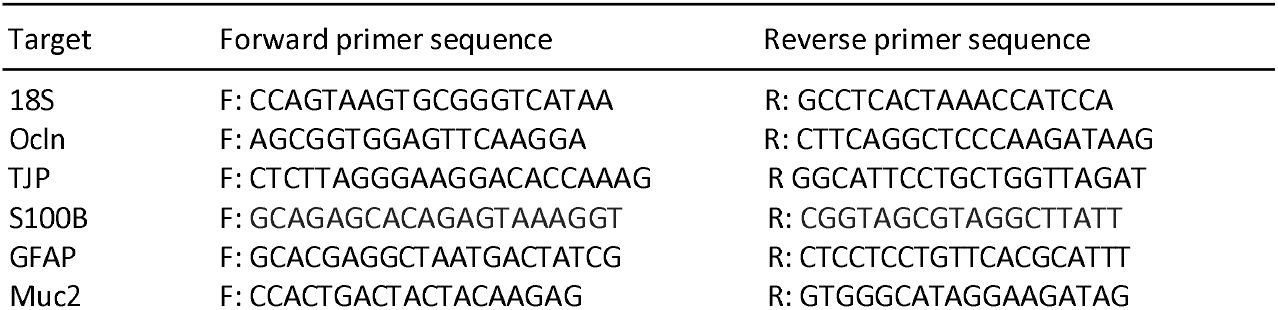
Primers used for qPCR and PCR

### Microbiome analysis

Microbiome samples underwent 16S rRNA sequencing to identify bacterial genera/species. Standard protocols from earthmicrobiome.org were used for library preparation. DNeasy Powersoil Kit (Qiagen, Germantown, MD) was used to extract DNA and PCR was performed to amplify the V4 hypervariable region of the 16S rRNA gene with individually barcoded primers (515F-806R; Parada et al., 2016; Apprill et al., 2015; Caporaso et al., 2011, 2012). The PCR product was then purified using PCR Purification Kit (Qiagen). The concentration of DNA was measured using Quant-iT Picogreen Assay (Thermofisher Scientific) and an equimolar pool was made of all samples. The pool was sequenced at the MGH Next-Generation Sequencing Core using an Illumina MiSeq Instrument that performed 20,000 paired-end 250bp reads per sample (Illumina, San Diego, CA, USA).

The sequenced raw fastq files were processed with QIIME software package (v. 2018.2.0) (Bolyen et al., 2019). The sequences with low quality score (on average less than 25) were truncated to 240bp and by using deblur algorithm with default settings the erroneous sequence reads were filtered (Amir et al., 2017). The remaining high-quality sequences were aligned with mafft plugin (Katoh and Standley, 2013). Next, the aligned sequences were masked to remove highly variable positions and a phylogenetic tree was generated from the masked alignment by FastTree plugin (Price et al., 2010). Alpha (richness and evenness) and beta diversity metrics (weighted and unweighted UniFrac, Bray Curtis and Jaccard dissimilarity) and Principal Coordinate Analysis plots were generated by default QIIME2 plugins (Bolyen et al., 2019; Lozupone and Knight, 2005; Vázquez-Baeza et al., 2013). To assign taxonomies to our sequences we used QIIME2’s feature-classifier plugin and pre-trained Naïve Bayes classifier, which has been trained on the Greengenes 13_8 99% operational taxonomic units (OTUs). Differential abundance analysis of OTUs were performed by ANCOM (Mandal et al., 2015), where significant differences were indicated by the W value. The extent of differences in the taxonomic profiles was substantiated further by analyzing LDA effect sizes calculated through LEfSe (LDA >4, P < 0.05) (Segata et al., 2011).

We used 16S ribosomal RNA sequencing to assess the microbial composition of the cecal microbiome at PND 60 in male and female prairie vole offspring born via VD or CS and cross-fostered to surrogate dams at birth. An individual’s microbial community structure can be assessed by alpha diversity which includes measures of richness (the number of different bacterial species present) and evenness (whether all species have a similar abundance within the community). Gut microbes reside in the mucus layer of the gut lumen adjacent to the intestinal epithelium and can modulate the development of the intestinal wall and villi, enteric glia that reside within the mucosal wall, and tight junction (TJ) proteins that regulate pore formation between epithelial cells. These TJ proteins control the movement of bacteria and food particles across the epithelial surface and their dysregulation can cause “gut leakiness” and stimulate inflammation. We therefore measured the expression of five TJ genes.

### Analysis

For measures involving repeated measures (USVs, thermography) results were analyzed using the lmer() function from the *Ime4* library (Bates et al., 2015, p. 4) followed by calling the anova() function on the resulting model. Otherwise, results were analyzed using ANOVA. In both cases, sex and birth mode were considered as main effects. For the pup measures of USV production and themography, a repeated measures linear mixed effects model with time (age in the case of USVs, minutes of test in the case of thermography) and birth mode (CS vs. VD) as main effects and accounting for weight and litter as random effects. Autoradiographic analyses of OTR and V1aR densities were compared on a regional basis, first by bilaterally averaging each subject’s measures, then using a repeated measures linear mixed effects model, so as to accommodate the repeated sampling of each brain region (two slices, often with bilateral representation). Outlying autoradiographic data points were detected and flagged automatically using Grubb’s test and then manually confirmed. Post-hoc analyses were carried out using Tukey’s HSD. When results were not normally distributed, data were analyzed by Kruskal-Wallis and Mann-Whitney tests. Data are reported as means plus/minus standard error, alpha = 0.05, and effect sizes reported as either η^2^ or Cohen’s *d.*

## RESULTS

### Weights, Experiment 1

VD and CS litters did not differ in litter size; VD litters had 4.52 ± 0.28 pups, and CS litters had 4.52 ± 0.25 pups. At delivery/discovery, VD pups (n = 95) weighed more than CS pups (n = 95, p = 0.001, *d* = 1.15); VD pups weighed 2.81 ± 0.08g, and CS pups weighed 2.41 ± 0.07g. On PND-1, VD pups weighed 3.15 ± 0.04g, and CS pups weighed 2.90 ± 0.04g (p > 0.05); on PND-4, VD pups weighed 4.37 ± 0.07g, and CS pups weighed 4.23 ± 0.07g (p > 0.05); on PND-7, VD litters weighed 25.38 ± 1.2g and CS litters weighed 23.72 ± 1.19g (p > 0.05).

### Weights, Experiment 2

Again, there was no significant difference in litter size; VD litters had 5 ± 0.7 pups, CS litters had 4.38 ± 0.4 pups, and CS+OXT litters had 3.71 ± 0.5 pups. At delivery/discovery, there was a significant effect of birth mode on pup weight (F(2,20) = 9.071, p = 0.002), with CS pups weighing less than pups of either other group (p < 0.004 for both comparisons, η^2^ = 0.48); VD pups weighed 2.85 ± 0.11g (n = 34), CS pups weighed 2.39 ± 0.08g (n = 33), and CS+OXT pups weighed 2.88 ± 0.09g (n = 28). On PND-1 there was a significant main effect of birth mode on pup weight, with both VD and CS+OXT pups weighing more than CS pups (p = 0.019 and p = 0.014 respectively, η^2^ = 0.28). However, by PND-4, there was no significant difference between groups’ pup weights. We were not able to make conclusions about litter weights on PND-7 due to our failure to preserve body weight measures.

### USVs, Experiment 1

Across PND-1 and PND-4, 288 recordings were made from pups (68 CS pups and 88 VD pups) from 43 litters (21 CS litters and 22 VD litters). In analyzing the total number of USVs, there were significant main effects of birth mode (F(1,21.08) = 4.65, p = 0.04, η^2^ = 0.18), age (F(1,169.19) = 14.5, p < 0.001, η^2^ = 0.08), and weight (F(1,85.3) = 5.86, p = 0.017, η^2^ = 0.06), along with a birth mode by age interaction effect (F(1,230) = 4.35, p = 0.038, η^2^ = 0.02), which post hoc analysis revealed as VD pups producing significantly fewer USVs on PND-4 compared to PND-1 (p = 0.002, Figure 1A) and CS pups producing more USVs than VD pups on PND-4 (p = 0.018). CS pups showed no age-related decline in USVs (p = 0.643). There was no effect of sex.

### USVs, Experiment 2

Across PND-1 and PND-4, 172 recordings were made from pups (32 CS pups, 33 VD pups, and 25 CS+OXT pups) of 23 litters (8 CS litters, 8 VD litters, and 7 CS+OXT litters). In analyzing the total number of USVs, there was a birth mode by age interaction effect (F(2,154.54) = 3.67, p = 0.028, η^2^ = 0.05), however no post-hoc comparisons were significant. There was no effect of sex.

### Thermography, Experiment 1

A total of 19 CS and 21 VD litters were analyzed. Litters of both birth mode conditions experienced a decline in surface warmth throughout testing, but CS litters consistently emitted less warmth (Figure 2). There was an expected main effect of time (F(5,175) = 268.47, p < 0.001, η^2^ = 0.88), as well as a main effect of birth mode (F(1,34) = 5.57, p = 0.024, η^2^ = 0.14) and an interaction between time and birth mode (F(5,175) = 2.38, p = 0.04, η^2^ = 0.06). Post hoc analyses revealed CS litters to have less surface area > 30.5°C during the three initial warm phases of testing even after controlling for litter size and total litter weight (minutes 0-30, p < 0.025 for all comparisons). For huddling behavior, there were also main effects of both time (F(5,146.63) = 13.14, p < 0.001, η2 = 0.31) and birth mode (F(1,150.3) = 6.55, p = 0.011, η2 = 0.04), along with a time by birth mode interaction (F(5,146.63) = 2.88, p = 0.016, η2 = 0.09), which post hoc analysis revealed as CS litters having significantly more disparate huddles during the first 10 minutes of testing (p < 0.001). For the total perimeter of a huddled litter, there were main effects of litter weight (F(1,137.79) = 11.48, p < 0.001, η2 = 0.08), time (F(1,137.79) = 11.48, p < 0.001, η2 = 0.38), and birth mode (F(1,135.46) = 26.07, p < 0.001, η2 = 0.16), which post hoc analysis revealed as CS litters displaying higher total perimeters of their huddles during the first twenty minutes (p < 0.041 for phases 1,2, 4 and 6).

### Consolation, Experiment 1

A total of 12 CS males, 5 CS females, 8 VD males and 8 VD females were tested. There was no effect of either sex or birth mode on any consolation behavior under observation (p > 0.05 for all comparisons). Anecdotally, both VD and CS animals displayed the expected set of conciliatory behaviors directed at their stressed cage-mates (i.e. investigation and allogrooming).

### Open Field, Experiment 1

A total of 16 CS males, 7 CS females, 17 VD males and 20 VD females were tested. There was no effect of sex or birth mode on time in the center of the OFT (p > 0.05 for both comparisons). However, there was a sex by birth mode interaction on total distance traveled (F(1,56) = 7.096, p = 0.01, η2 = 0.11), however no post-hoc comparisons were significant.

### Open Field, Experiment 2

A total of 13 CS males, 21 CS females, 10 VD males, 20 VD females, 11 CS+OXT males and 13 CS+OXT females were tested. Again, there was no effect of either sex or birth mode on time in the center of the OFT (p > 0.05 for both comparisons). There was however, a main effect of sex (F(2,79) = 4.096, p = 0.046, η2 = 0.05), with greater locomotion in males, and a trend toward an effect of birth mode on total distance traveled, (p = 0.057), such that the two CS conditions, when collapsed across Experiments 1 and 2, moved significantly more than VD counterparts (p = 0.016).

### Alloparenting, Experiments 1 and 2

A total of 17 CS males, 9 CS females, 20 VD males and 23 VD females were tested in Experiment 1 and in Experiment 2, 11 CS males, 18 CS females, 8 VD males, 19 VD females, 9 CS+OXT males and 13 CS+OXT females were tested. In Experiment 1, females attacked pups at a higher rate than males (chi-squared = 6.42, p = 0.011). Although we observed no treatment nor sex by treatment interactions in the proportion of animals attacking the pup, this still left us with a small sample size of female CS animals that displayed alloparental behavior (only 4 out of 9 CS females did not attack the pup). Because of this, the alloparenting results for Experiments 1 and 2 were combined into a single analysis. There was no effect of birth mode on any alloparental behavior under observation (Figure 3, p > 0.05 for all comparisons). Females were generally less alloparental than males, displaying for instance less time in contact with the pup (F(1,103) = 12.762, p < 0.001, η2 = 0.02) and less time huddling the pup (F(1,103) = 8.907, p = 0.004, η2 = 0.08). These same patterns persisted when we analyzed Experiments 1 and 2 separately (data not shown).

### Partner Preference, Experiment 1

A total of 17 CS males, 11 CS females, 22 VD males, and 23 VD females were tested. The setup of the Partner Preference Test is shown in Figure 4A. Both male and female VD animals formed partner preferences as defined as spending significantly more time in close social contact with the Partner vs. the Stranger (p < 0.05 for both comparisons, Figure 4B). Neither male nor female CS animals formed partner preferences

### Partner Preference, Experiment 2

A total of 4 CS males, 13 CS females, 6 VD males, 9 VD females, 10 CS+OXT males and 10 CS+OXT females were tested. Again, both male and female VD animals formed partner preferences (p < 0.05 for both comparisons, Figure 4C) and neither male nor female CS animals formed partner preferences (p > 0.05 for both comparisons). Both male and female CS+OXT animals formed partner preferences (p < 0.05 for both comparisons). To examine the relationship between the length of gestation and partner preference formation in adulthood, we correlated birth weight with selectivity in the PPT, defined here as the difference between time spent with the Partner vs. the Stranger divided by the total time in close social contact. In neither Experiment 1 nor Experiment 2 was birth weight significantly correlated with selectivity in any condition.

### Microbiome, Experiment 1

A total of 10 CS males, 10 VD males, 9 CS females and 10 VD females were tested. We did not observe any significant differences in richness or evenness for males (observed OTUs: p=0.733; Fig. 5A; Pielou’s evenness: p=0.762; Fig. 5B; Faith’s phylogenetic diversity: p=0.364; Shannon: p=1.00) or females (observed OTUs: p=0.221; Fig. 5C; Pielou’s evenness: p=0.514; Fig. 5D; Faith’s phylogenetic diversity: p=0.624; Shannon: p=0.462) based on birth mode. Beta diversity refers to a difference in microbial community structure between individuals. Principle Coordinate Analysis (PCoA) of beta diversity indices revealed no significant differences in the clustering of microbiome profiles between CS and VD animals in either males g (Jaccard dissimilarity: p=0.056; Bray-Curtis dissimilarity: p=0.068: Fig. 5E) or females (Jaccard dissimilarity: p=0.123; Bray-Curtis dissimilarity: p=0.607; Fig. 5E). Other beta indices revealed no significant differences for males (unweighted-unifrac dissimilarity: p=0.180; weighted-unifrac dissimilarity: p=0.333) or females (unweighted-unifrac dissimilarity: p=0.461; weighted-unifrac dissimilarity: p=0.366). Differences were not found at the level of individual taxa using ANCOM testing. Linear discriminant analysis effect size (LEfSe) revealed differentially abundant bacteria in male VD and CS offspring. At the order level, Clostridiales was more abundant in CS males than VD males (Fig. 6A), at family level, Prevotellaceae was more abundant in VD males than CS males and (Fig. 6B) and at the genus level, Ruminoococcus was more abundant in CS males than VD males (Fig. 6C).

**Figure 5.**
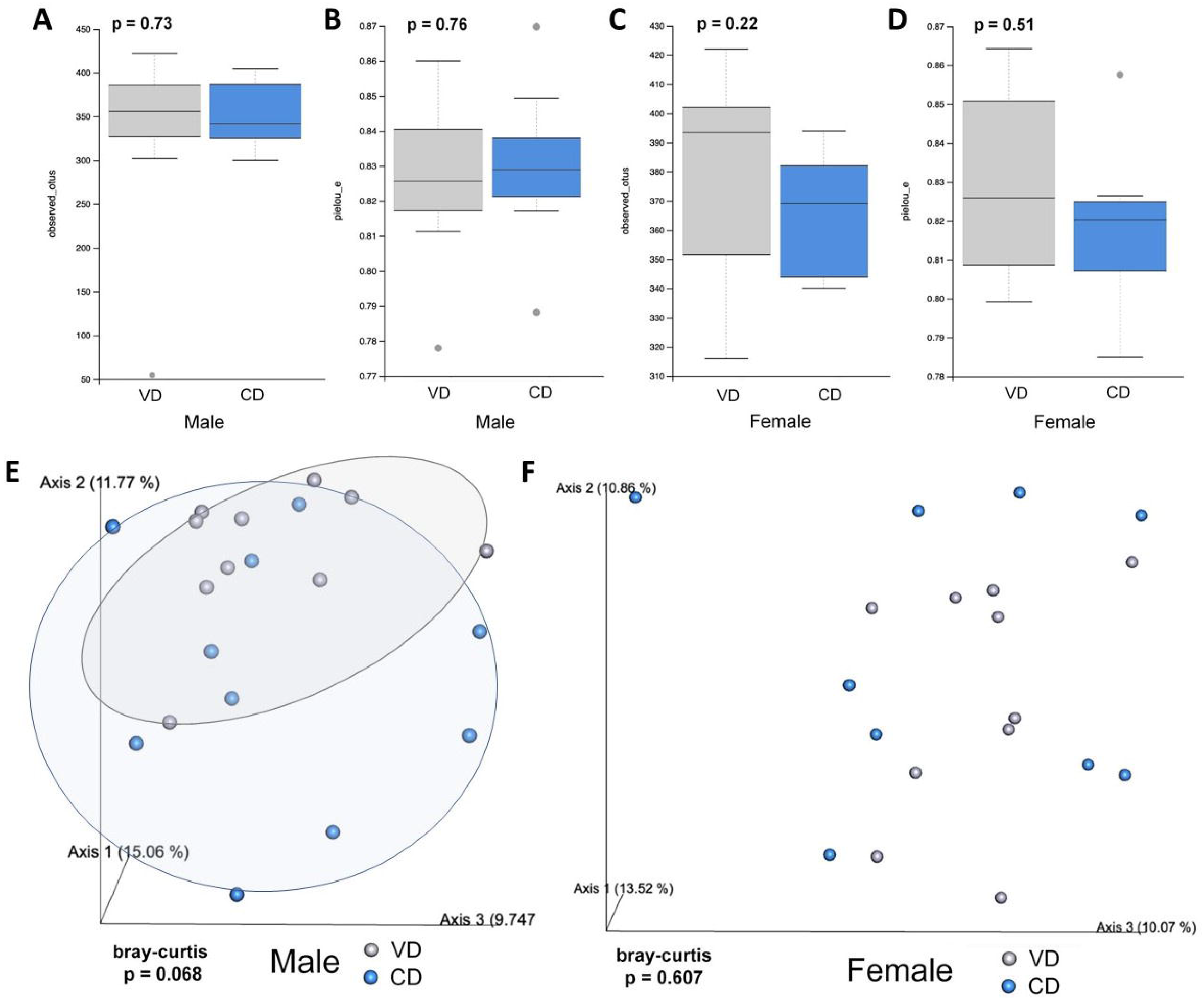
CS dose not shift the diversity or overall composition of the gut microbiome as compared to VD using metrics of alpha and beta diversity. **A-D**, Observed OTUs and Pielou’s evenness were not significantly different in CS males as compared to VD males (A, OTUs, N=10/group, p=0.73; B, Pielou’s evenness N=10/group, p=0.76) or in CS females as compared to VD females (C, OTUs, N=10/group, p=0.22; D, Pielou’s evenness N=9-10/group, p=0.51). **E-F**, There were no significant differences in the clustering of gut microbiota between CS and VD animals for either males (E, N=10/group, Bray-curtis dissimilarity, p=0.068; N=10/group, Jaccard dissimilarity, p=0.056, data not shown) or females (F, N=9-10/ group, Bray-Curtis dissimilarity, p=0.607; N=9-10/group, Jaccard dissimilarity, p=0.123, data not shown).

**Figure 6.**
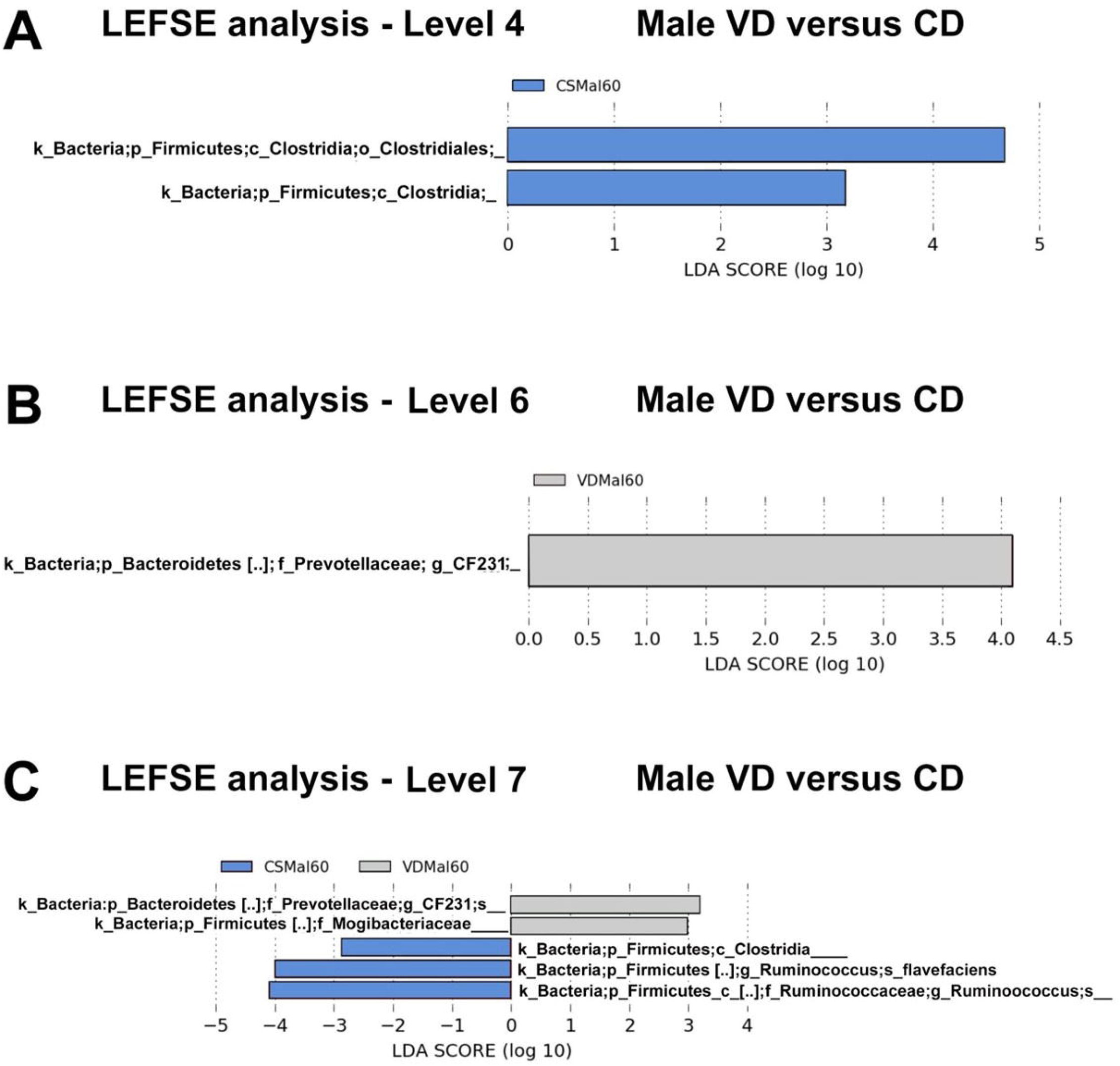
Linear discriminant analysis effect size (LEfSe) indicated differentially abundant bacteria in male VD and CS offspring at the order (A), family (B) and genus (C) levels. **A**, Clostridiales was more abundant in CS males than VD males. **B**, Prevotellaceae was more abundant in VD males than CS males. **C**, Ruminoococcus was more abundant in CS males than VD males. LDA >4, P < 0.05.

### Gut Gene expression, Experiment 1

Within the terminal ileum, a significant decrease was observed in the expression of occludin (OCLN) but not tight-junction protein (TJP; also known as zonula occludens-1 or ZO-1), two TJ scaffolding proteins important for barrier stability (Fig. 7A). There were also no differences in the expression of S100 calcium-binding protein B (S100B) and glial fibrillary acidic protein (GFAP), well-known markers of mucosal enteric glia (Fig. 7A). In contrast, within the proximal colon, the expression of OCLN and TJP were significantly decreased in CS male offspring as compared to VD male offspring (Fig. 7B). In addition, the expression of enteric glial markers S100B and GFAP were significantly decreased in CS males as compared to VD males (Fig. 7B), while no difference in mucin 2 (MUC2), a scaffolding protein that forms the mucus layer, were observed in either males or females (Fig. 7B). These differences in gene expression in CS males were large in effect size (*d* > 1).

**Figure 7.**
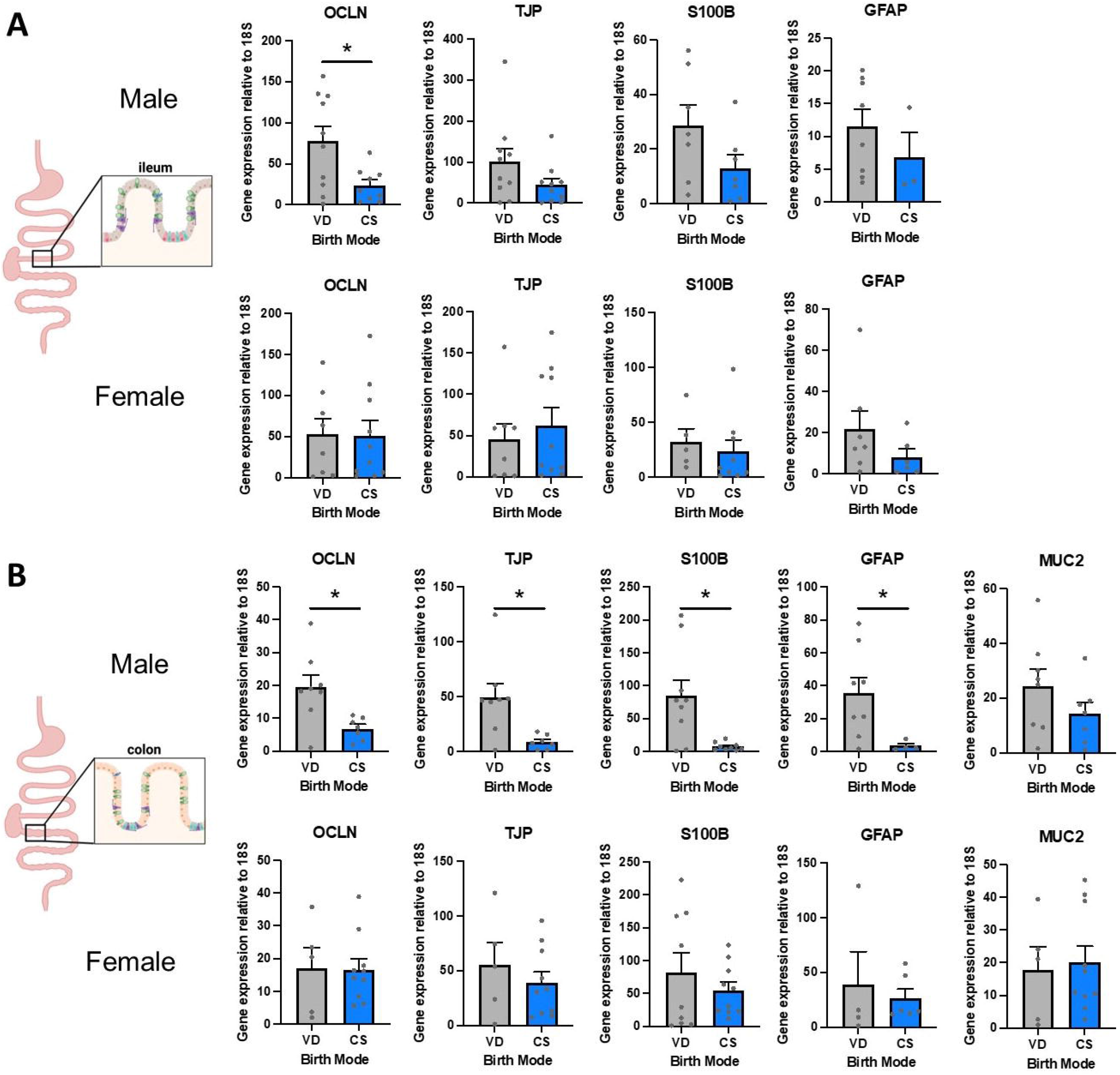
Significant differences in the gene expression of tight junction proteins (TJs) and enteric glial markers within the colon of CS versus VD males. **A**, Schematic representing the section of the terminal ileum assessed and the quantification of gene expression for TJ proteins occludin (OCLN) and tight junction protein-1 (TJP) and enteric glia markers S100 calcium-binding protein B (S100B) and glial fibrillary acidic protein (GFAP). No significant differences were observed in the ileum of CS versus VD males except for OCLN (N=3-10/group, unpaired t-test; OCLN: treatment: p=0.018, TJP: treatment: p=0.123, S100B: treatment: p=0.109 and GFAP: treatment: p=0.349). No significant differences were observed for CS versus VD females (N=5-10/group, unpaired t-test; OCLN: treatment: p=0.942, TJP: treatment: p=0.567, S100B: treatment: p=0.600 and GFAP: treatment: p=0.218). **B**, Schematic representing the section of the proximal colon assessed and the quantification of gene expression for TJ proteins OCLN and TJP, enteric glia markers S100B and GFAP, and mucin 2 (MUC2), a major component of intestinal mucus. There was a significant decrease in OCLN, TJP, S100B and GFAP in CS males compared to VD males (N=4-9/group, unpaired t-test; OCLN: treatment: p=0.012, TJP: treatment: p=0.011, S100B: treatment: p=0.006 and GFAP: treatment: p=0.046) while no differences were observed for females based on birth mode (N=4-10/group, unpaired t-test; OCLN: treatment: p=0.944, TJP: treatment: p=0.443, S100B: treatment: p=0.392 and GFAP: treatment: p=0.654). Data represent mean ± SEM, *p<0.05. CS: cesarean section, VD: vaginal delivery. No differences were observed for MUC2 based on birth mode for males or females (males, N=7-8/group, unpaired t-test; MUC2: treatment: p=0.208; females, N=5-10/group, unpaired t-test; MUC2: treatment: p=0.777).

### Autoradiography, Experiment 1

Overall, there were few differences in either OTR or V1aR observed. In the agranular insular cortex, we noted a main effect of birth mode on OTR density (F(1,46) = 4.375, p = 0.042, η2 = 0.08), with CS animals having greater OTR. In the claustrum, there was a main effect of birth mode on OTR density (F(1,48) = 6.827, p = 0.012, η2 = 0.18), with CS animals having greater OTR. However, in no other brain region did we note an effect of birth mode, and therefore, considering the number of comparisons made, we cannot place much confidence in these two differences.

In the central amygdala, we observed a sex by birth mode interaction on OTR (F(1,46) = 5.573, p = 0.0225, η2 = 0.1) such that CS females had greater OTR than VD females (p = 0.023). Across the lateral septum at 2 anterior-posterior positions, there were consistent main effects of sex on OTR (F(1,48) = 5.111, p = 0.028, η2 = 0.09 and F(1,48) = 4.63, p = 0.036, η2 = 0.08), with females having greater OTR. In the cortical amygdala, we noted a main effect of sex on V1aR (F(1,45) = 4.616, p = 0.037, η2 = 0.08), with males having greater V1aR density. In the ventral pallidum, there was a main effect of sex (F(1,45) = 4.051, p = 0.05, η2 = 0.06), such that females had greater V1aR density, as well as a sex by birth mode interaction (F(1,45) = 6.6107, p = 0.014, η2 = 0.06), however no post-hoc comparisons of relevance were significant.

## DISCUSSION

In this study, we observed a failure of partner preference formation in voles delivered by CS, which we were able to replicate in a follow-up study; this effect was prevented using direct OXT administration to the pup. Briefly, in Experiment 1, CS pups were found to produce more USVs on PND-4, had lower skin temperature and huddled less cohesively on PND-7, failed to form partner preferences as adults, and showed minor changes in OTR density in the agranular insular cortex and claustrum. CS pups in Experiment 2 tended to show the opposite pattern in terms of USVs, with CS pups producing fewer calls; unfortunately, we were not able to complete thermographic analysis at PND-7, nor autoradiographic or metabolomic analyses upon sacrifice. The underlying mechanism of the observed behavioral differences awaits further investigation because neither of the two candidates investigated here, neuropeptide receptor density within the brain and microbiome diversity in the gut, offered a robust explanation. A single brain region (the agranular insular cortex) was found to be affected by CS delivery in terms of OXTR density in adulthood. Similarly, the most noteworthy finding of the present study is therefore that voles delivered via CS fail to form partner preferences, an effect which can be prevented by treating CS pups with 0.25 mg/kg OXT immediately after CS delivery.

Among other *tocosematic* or ‘birth-signaling’ hormones, OXT is an important factor for coordinating the transition to extrauterine life (Kenkel, 2020; Kenkel et al., 2014). Both mother and neonate release OXT during delivery, with each needing to accomplish tremendous physiological changes. Disruptions of OXT signaling at delivery can affect a broad range of offspring outcomes in the acute phase, including neuroinflammation (Kingsbury and Bilbo, 2019), pain sensitivity (Mazzuca et al., 2011), and feeding behavior (Schaller et al., 2010). Still more long-term offspring neurobiological outcomes may be influenced by perinatal OXT manipulations, including changes in the expression of: OXT and its receptor (Kenkel et al., 2019b; Yamamoto et al., 2004), vasopressin and its receptor (Bales et al., 2007a; Yamamoto et al., 2004), estrogen receptor alpha (Kramer et al., 2007) serotonin and dopamine (Eaton et al., 2012; Hashemi et al., 2013) within the brain. Moreover, delivery by CS has been shown to alter the balance of dopamine signaling in adulthood in rats (Boksa and Zhang, 2008; El-Khodor and Boksa, 2001, 1997) and dopamine is a critical regulator of pairbonding (Aragona and Wang, 2009). Because birth is a sensitive period in terms of the long-term neurobiological effects of many birth-signaling hormones, there are thus multiple potential neuroendocrine routes through which CS could have altered social bonding.

Previous work in mice has found that CS pups either produce more calls on PND-9 (Chiesa et al., 2020, 2019; Morais et al., 2020) or produce an equivalent number of calls (Castillo-Ruiz et al., 2018; Morais et al., 2021) or USVs with decreased amplitudes (Castillo-Ruiz et al., 2018) compared to VD counterparts, however, these studies were all in mice at a later stage of development than the current work. Our prior study of VD vole pups on PNDs 1 and 4 found that exposure to prenatal OXT led to increased USV production (Kenkel et al., 2019b). This led us to predict that CS would diminish calls, which we observed in Experiment 2, but the results of Experiment 1 revealed the opposite pattern. In the same prior study on prenatal OXT administration, we also observed that OXT led to a broadly gregarious phenotype in adult offspring. OXT-exposed voles displayed increased alloparental responsiveness (in two separate experiments) and increased time in close social contact with opposite-sex adults (in the only experiment that addressed this behavior). We did not observe an impact of birth mode on alloparental behavior in the present study. As has been observed previously in prairie voles, we did find males to be more alloparental than females (Kenkel et al., 2016). Importantly, prenatal OXT exposure did not affect the selectivity aspect of pair-bonding in that prior work, i.e. voles were no more nor less monogamous, whereas in the present study, CS impacted the selectivity of pairbonding such that CS animals failed to form selective partner preferences. We are unaware of any work that has considered pair-bonding or romantic attachments in humans as a function of birth mode.

Direct treatment of CS pups with 0.25 mg/kg OXT immediately after delivery was able to prevent the deficit in pair-bonding observed in CS offspring. This is in line with our hypothesis that CS can affect neurodevelopment by impacting levels of the birth-signaling hormones during the sensitive period around birth. This also matches recently published work which found that daily OXT treatment (~0.1 or 1 mg/kg) between days 1 and 5 could rescue social behavior deficits in mice delivered by CS (Morais et al., 2021). In that study, CS pups did not experience different levels of parental care -an important consideration that we hope to pursue in the future. While we observed no effect of CS on anxiety-like behavior in the OFT in the present study, the recent study by Morais and colleagues found that CS led to diminished marble burying behavior in offspring mice, which could be prevented by OXT treatment (Morais et al., 2021). CS mice also showed less preference for their natal nest, which was restored by treatment with the ~1 mg/kg OXT dose. In that same study, CS also led to decreased locomotion in an aversive open field, which was not rescued by OXT treatment. Finally, CS mice showed deficits in social novelty preference, which were restored by the ~0.1 mg/kg OXT dose. CS delivery affects the levels of many other hormones beyond merely OXT (Kenkel, 2020), but the pattern of results from the present study and that of Morais and colleagues points to exogenous OXT being able to avert the neurodevelopmental consequences of CS delivery, likely by effectively normalizing OXT levels during the sensitive perinatal period. While OXT appears important for the transition to independent homeostasis in many mammals (Kenkel et al., 2014), there may be species differences in the extent to which CS affects OXT levels in the neonate. In humans, CS is associated with a 37-75% reduction in OXT levels (Chard et al., 1971; de Geest et al., 1985; Kuwabara et al., 1987; Marchini et al., 1988). We do not currently know the extent to which CS impacts OXT levels in neonatal prairie voles.

One important alternative explanation to the present findings which must be considered is that offspring in the CS condition were affected by early delivery and low birth weight. In Experiments 1 and 2 CS pups weighed less than VD pups. At delivery, CS pups weighed ~2.4g while VD pups weighed ~2.8g. Some of this difference in weight must be attributed to VD pups having 0-12 hours of nursing prior to discovery. CS negatively impacts feeding reflexes in human neonates (Heidarzadeh et al., 2016) and newborn rodents (Alberts and Ronca, 2012). However, we cannot completely discount the possible role for early delivery on the developmental outcomes observed here. A meta-analysis of 4.4 million people found that adults who were born preterm or of low birth weight were less likely to ever experience a romantic relationship, sexual intercourse, or parenthood than peers born full-term (odds ratios of 0.72, 0.43, and 0.77 respectively) (Mendonça et al., 2019). There are clearly many differences between preterm and CS delivery, but this does establish that birth experience can shape social bonding in adulthood. Furthermore, OXT is not the only birth-signaling hormone affected by CS. In rats, CS delivery influences the regulation of dopamine in later life via diminished levels of the birth signaling hormone epinephrine (Boksa and Zhang, 2008; El-Khodor and Boksa, 2001, 1997), which would also affect social attachment as dopamine is critical to pair-bond formation and maintenance in prairie voles (Aragona et al., 2003). It stands to reason that treatment with vasopressin, corticosterone, epinephrine or norepinephrine could each also normalize development in CS offspring.

Similar to prior work examining the paraventricular nucleus of the hypothalamus in mice (Morais et al., 2021), we did not find evidence for substantial changes in OTR and V1aR in the adult brains of CS offspring. Although we observed CS animals having greater OTR in the agranular insular cortex and claustrum, as well as CS females having greater OTR in the central amygdala, we must consider the large number of comparisons made. For the time being, these findings are promising directions for future work to either confirm or reject. One intriguing possibility comes from recent work indicating that ligand expression patterns rather than those of the receptors may be impacted by CS delivery (Ramlall et al., 2021). Ramlall and colleagues recently reported that CS delivery led to fewer and smaller vasopressin cells and smaller OXT cells in the paraventricular nucleus of adult mice (Ramlall et al., 2021). Diminished vasopressin or OXT signaling could certainly lead to deficits in pair-bonding in prairie voles (Cho et al., 1999).

Overall, our microbiome analyses did not yield any significant differences in alpha diversity metrics or beta diversity metrics between CS versus VD adult prairie vole offspring that were cross-fostered at birth, consistent with what has been observed in mice for measures of alpha diversity and beta diversity using principal coordinate analyses (Morais et al., 2021). We did observe significant changes in overall abundance of bacterial taxa at the level of order, family and genus, similar to (Morias et al., 2020). Within the present study, Clostridiales, Prevotellaceae, and Ruminoococcus in adult prairie vole offspring were differentially affected by birth mode. Changes in Ruminoococcus at the genus level were also shown to be impacted by birth mode in mice (Morais et al., 2021). It is becoming increasingly recognized that there is an intricate interplay between gut microbes and the maturation of the gastrointestinal system, including the development of gut structure and immune cell components (Kabouridis et al., 2015; Park et al., 2016; Peck et al., 2017). To our knowledge, we are the first to show an alteration in the expression of intestinal TJ proteins and enteric glia markers in gut tissue of rodent offspring delivered by CS as compared to VD. Morias et al 2021 noted that mice delivered via CS had increased gastrointestinal motility compared to VD mice, an effect that could be reversed with OT treatment (Morais et al., 2021). Our findings of dramatically decreased expression of enteric glial markers in male CS offspring suggest that enteric glia development and function may be significantly altered by birth mode, with sex-dependent effects. This finding requires follow-up as enteric glia are important regulators of gut motility, mucosal homeostasis and immunity, and gut barrier function (Grubišić and Gulbransen, 2017). Concurrent with the changes in enteric glia markers, we observed decreased expression of TJ proteins OCLN and TJP (ZO-1) in male CS offspring compared to VD offspring. Similarly, a greater change in the expression of TJ proteins and permeability following DSS-induced colitis was observed in male mice compared to female mice within the zonulin transgenic mouse model (Miranda-Ribera et al., 2019, p.). Given the abundance and diversity of TJ proteins, it is not currently clear if the expression changes in OCLN and TJP we observed here impact gut permeability and Morias et al. found that gastrointestinal permeability was not differentially impacted in CS and VD mouse offspring, unlike gastrointestinal transit (Morais et al., 2021). How birth mode impacts the dynamic interplay between gut microbes, intestinal development, and mucosal immunity requires further investigation.

Previous work in humans (Christensson et al., 1993) and lambs (Clarke et al., 1998, 1997) has found that CS impacts thermoregulation in offspring, which we take as evidence that the initiation of independent homeostasis has been compromised. Our present findings confirm this effect substantially further into postnatal development compared to previous work, which focused on the first postnatal day. Here, we observed lower surface temperatures in CS pups along with less cohesive huddling behavior. It remains for future work to examine how these differences relate; that is, are they independent or does less huddling lead to lower surface temperatures (or vice versa)? OXT is a potent thermoregulatory hormone, one which directly activates brown adipose tissue (Harshaw et al., 2018; Kasahara et al., 2013), thus by diminishing OXT levels during the sensitive period around delivery, CS could impact the onset of independent thermoregulation. Differences in thermoregulation would also contribute to the production of isolation-induced USVs, as temperature is arguably the largest driving factor for USV production (Kenkel et al., 2015). However, if a thermoregulatory difference persisted into adulthood, we would also have expected this to have impacted alloparenting, which we did not observe here. Alloparenting in prairie voles is heavily dependent on sensing the thermoregulatory state of the pup (Kenkel et al., 2015). In any case, changes in thermoregulation should be considered a possible avenue for how CS impacts development alongside neuroendocrine and microbial mechanisms. One major advantage of working with prairie voles is that they are adapted to cool climates and appear to avoid the chronic cold stress that mice and rats face in conventional ‘room temperature’ (22°) housing (Kenkel et al., 2021). This is especially important when considering the developmental consequences of cold stress on vulnerable pups, where prairie voles have the added benefit of biparental care providing additional warmth in the nest. Thus, we view the prairie vole as an especially well-suited model for studying the neurodevelopmental consequences of CS delivery.

This study should be viewed as an initial foray into the neurodevelopmental consequences of CS delivery, and it certainly not without limitations. For instance, in the consolation test, sibling cage mates were all of the same experimental condition. Thus, a change in conciliatory responsiveness on the part of the subject may have been masked by a complementary change in distress signaling on the part of the cage mate. Moreover, sample sizes were limited and somewhat inconsistent. Sample sizes were most robust for USV measures, however findings there were contradictory. In other tests, limited human resources prevented comprehensive testing of all subjects.

CS is worthy of further investigation given its epidemiological associations with various later life health outcomes and also simply due to its prevalence. Regardless of whether CS delivery is causally related to immune or metabolic or social differences, CS affects the signaling of several important hormones during a sensitive period in development, which raises the prospect of enduring consequences. While the cost/benefits of CS have traditionally focused on the acute term near birth, studies of longer-term effects are especially needed given alterations in the nervous and gastrointestinal systems and behavior shown here and related studies (Morias, 2020, 2021). These consequences may or may not rise to the level of a clinical diagnosis, but deserve further investigation regardless. Given that CS rates are rising around the world and stand already at nearly a third of all births in the U.S., we owe public health a fuller accounting of the benefits and drawbacks that CS delivery affords future generations and how therapeutic interventions during or after birth may mitigate some of the drawbacks.

---

Supplemental Figure 1

The analysis pipeline for autoradiographic histology. Raw images were first screened for tissues tears and folds, which were digitally removes from the analysis. Three images of each subject, matched for anterior-posterior position, were then registered manually registered to a prairie vole brain atlas (Yee et al., 2016). using Photoshop (Adobe Systems Inc., San Jose, CA). Images were then consolidated into group composites for the purpose of visualizing group differences. Heat maps were generated in Matlab for each group-by-group comparison by subtracting one group composite from another.

Supplemental Figure 2

Composite OTR autoradiographs are shown for each sex and birth mode combination across the three anterior-posterior levels.

Supplemental Figure 3

Composite V1aR autoradiographs are shown for each sex and birth mode combination across the three anterior-posterior levels.

Supplemental Table 1

Autoradiography results for OTR across all brain regions examined. * denotes a main effect of birth mode, § denotes a main effect of sex, and † denotes a birth mode by sex interaction.

Supplemental Table 1

Autoradiography results for V1aR across all brain regions examined. § denotes a main effect of sex.

## Acknowledgements

This work was funded by a grant from the Eunice Kennedy Shriver National Institute of Child Health and Development (NICHD), P01HD075750 and R21HD098603. The authors would also like to thank: the assistance of the animal care staff at Indiana University; Alexis Daughhetee, Rebecca Gray, Cynthia Stanton, and Nichol Crose for their hard work assisting with the behavioral experiments; Jennifer Ihedioha for technical assistance with gut PCR and Danielle Rendina for help with Lefse analyses; and the Center for the Integrative Study of Animal Behavior (CISAB) core facility at Indiana University, including David Sinkiewicz. Original data are available upon request.

