## Supplementary figures and images for "Lasting consequences on physiology and social behavior following cesarean delivery in prairie voles"

### Supplemental Figure 1

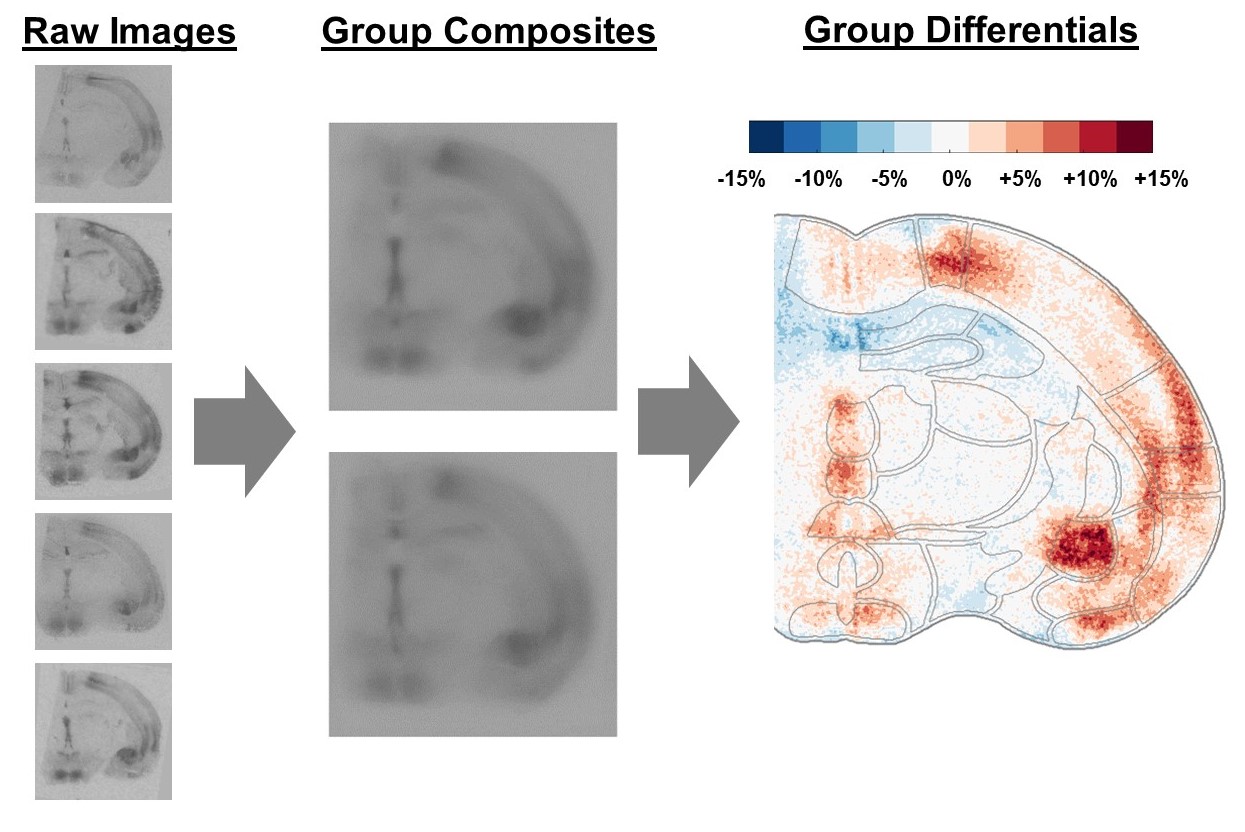

### Supplemental Figure 2

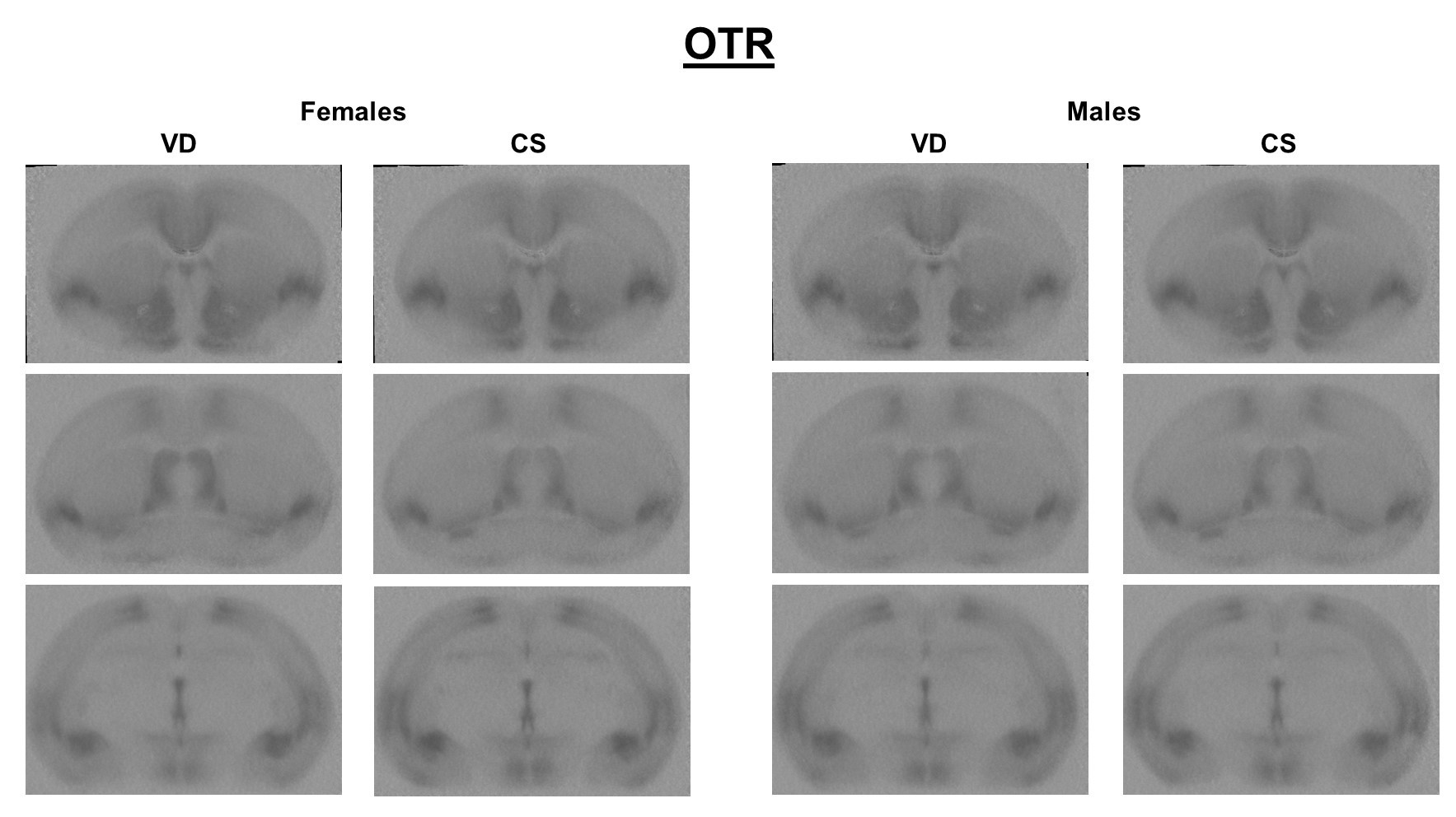

### Supplemental Figure 3

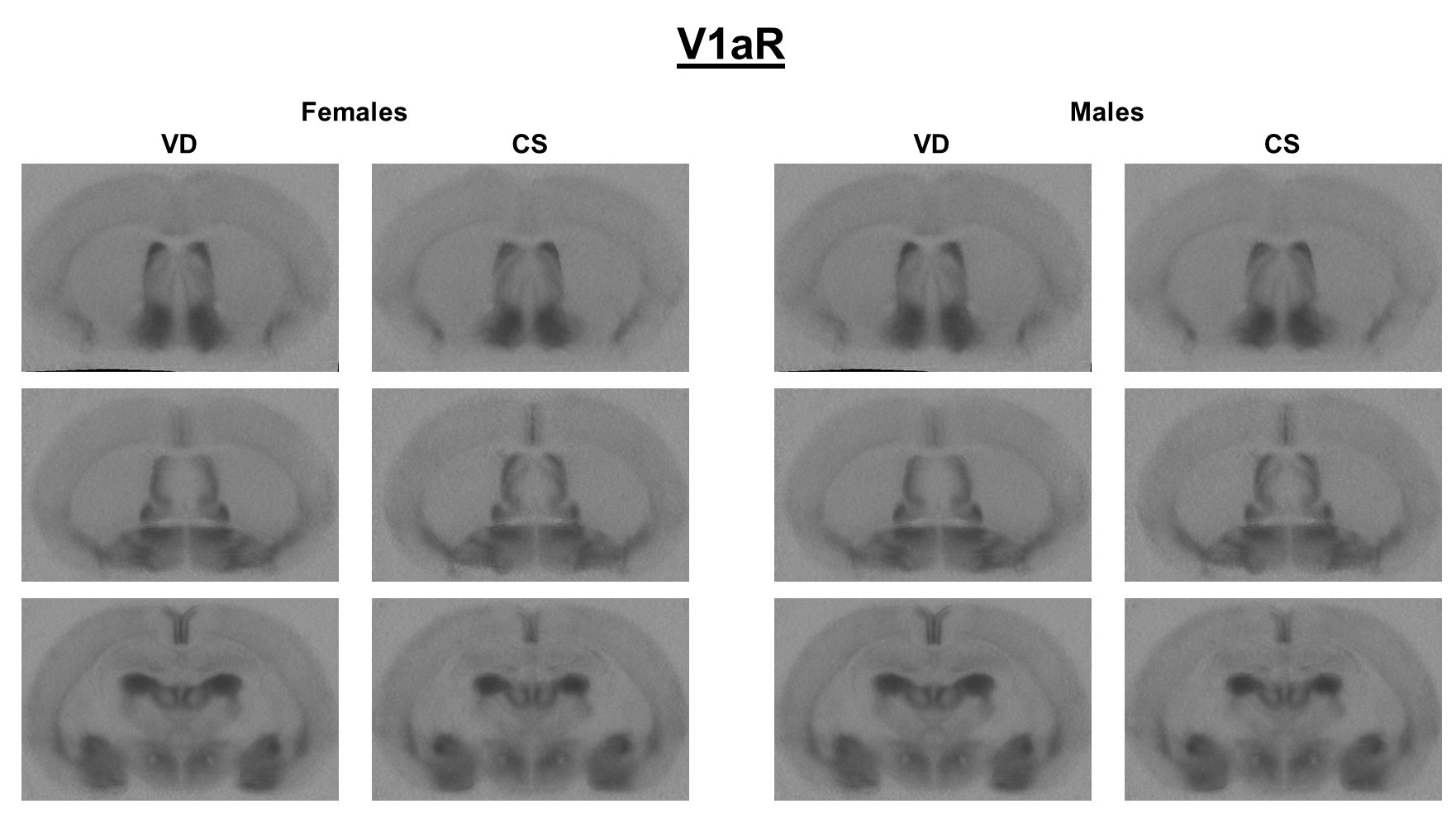
